# Projections from thalamic nucleus reuniens to medial septum enable extinction of remote fear memory

**DOI:** 10.1101/2024.05.20.594930

**Authors:** Kamil F. Tomaszewski, Magdalena Ziółkowska, Kacper Łukasiewicz, Anna Cały, Narges Sotoudeh, Monika Puchalska, Ahmad Salamian, Kasia Radwanska

**Author notes:** **Corresponding author:** Kasia Radwanska, Ph.D., Laboratory of Molecular Basis of Behavior, the Nencki Institute of Experimental Biology of Polish Academy of Sciences, 3 Pasteur St., Warsaw 02-093, Poland.

## Abstract

Aversive experiences lead to the formation of long-lasting memories. Despite the need to better understand how enduring fear memories can be attenuated, the underlying brain circuits remain largely unknown. In this study, employing a combination of genetic manipulations, neuronal circuit mapping, and chemogenetics in mice, we identify a new projection from the thalamic nucleus reuniens (RE) to the medial septum (MS), and show that this circuit is involved in the extinction of remote (30-day old), but not recent (1-day old), fear memories. These findings provide the first functional description of the RE→MS circuit and highlight the significance of the thalamo-septal regions in memory organization as a function of memory age, a phenomenon known as systems consolidation.

## Introduction

The distinction between dangerous and safe contexts, as well as the ability to update such information in a changing environment is crucial for animal survival, as mistakes can have costly consequences. Impairments in context-dependent behaviors have been associated with pathological conditions including post-traumatic stress disorder, schizophrenia, and substance abuse disorders ^1–4^. Therefore, understanding the molecular and cellular mechanisms that underlie the updating of contextual memories is vital for developing new therapeutic approaches^5–7^.

Contextual fear conditioning (CFC) represents a method to study contextual memories in laboratory animals ^4^. In this approach, a spatial context, the conditioned stimulus (CS), is repetitively paired with an aversive unconditioned stimulus (US), such as a mild electric shock. The observable freezing behavior serves as a measurable indicator of contextual fear memory. As exposure to the CS persists without the US, freezing behavior typically diminishes, accompanied by increased exploratory behavior. This process, known as fear extinction, unfolds as animals gradually recognize that the context no longer predicts impending shocks. To facilitate extinction, mice must actively retrieve and revise contextual memories, necessitating the presence of specific brain mechanisms. The “classic” neuronal circuit of contextual fear extinction includes medial prefrontal cortex (mPFC), amygdala, and hippocampus ^4,8^. However, the brain regions that contribute to memory recall may vary with memory age. In particular, memories for events initially depend upon the hippocampus, with time these memories become increasingly dependent upon the cortex for their expression ^9–11^. Post-learning communication between the hippocampus and cortex, via the thalamus, is thought to drive this time-dependent reorganization process called systems consolidation ^5,12–18^. This indicates that the extinction of remote memories may similarly rely on a re-distributed network, likely requiring additional players to the canonical extinction-mediating pathways of recent memories.

Calcium and calmodulin-dependent kinase II (CaMKII) is one of the key synaptic molecules that contributes not only to formation and updating of recent contextual memories but also to systems consolidation of remote memories ^15,19–23^. In particular, knock-in mice with a targeted Threonine 286 mutation to Alanine (T286A), that prevents the autophosphorylation of αCaMKII, have impaired formation and extinction of contextual memories ^20–25^, while heterozygous αCaMKII knock-out mice (αCaMKII^+/-^) have spared recent memory but severely impaired its long-term (>10 days) retention ^19^. Here, we tested the effect of heterozygous T286A mutation (T286A^+/-^) on extinction of recent and remote contextual fear memory. As T286A^+/-^ mutant mice showed impaired extinction of remote, but not recent, contextual fear memory, we used c-Fos immunostaining in this mouse model to study neuronal correlates of remote memory extinction. The analysis revealed a number of regions hyper- or hypoactivated in the heterozygotes during extinction of remote memory, including cortical, thalamic, hippocampal and amygdalar areas. While the contributions of hippocampal and cortical regions have been studied in detail, much less is known about how thalamic regions contribute to this memory reorganization process at different post-encoding delays. Accordingly, we focus here on the thalamic nucleus reuniens (RE) and medial septum (MS). We identified projections from anterior RE to MS (RE→MS) and showed that RE, MS and the RE→MS pathway are necessary for extinction of the remote, but not recent, contextual fear memory. Previous studies identified RE projections to CA1 area that are involved in extinction of recent contextual fear memory ^26–28^ and RE projections to the basolateral nucleus of the amygdala (BLA) to be involved in extinction of remote fear memories ^29^. Current results indicate a novel thalamo-septal pathway that is specifically engaged in the updating of remote memories. Our discoveries suggest that, akin to memory consolidation, the brain circuits involved in fear extinction undergo a spatial reorganization over time.

## Results

### Role of αCaMKII autophosphorylation in extinction of recent and remote contextual fear memory

We tested the effect of αCaMKII autophosphorylation of Threonine 286 on extinction of recent and remote contextual fear memory. To this end heterozygous autophosphorylation-deficent mutant mice (T286A^+/-^) and their wild-type (WT) littermates underwent contextual fear conditioning (CFC) with 5 electric shocks (unconditioned stimuli, US) in novel context. Next day, they were re-exposed to the training context for extinction of recent fear memories. At the beginning of the CFC freezing levels were low but increased during the training. At the beginning of the Recent Extinction session freezing levels were high in both experimental groups indicating contextual fear memory formation, and decreased within the session. Fear extinction memory was tested on the next day. Mice of both genotypes significantly decreased the frequency of freezing during the Test (T) as compared to the beginning of the Extinction session (E5) indicating formation of fear extinction memory. No significant differences in fear extinction memory was observed between the genotypes (**Figure 1A-C**).

**Figure 1.**
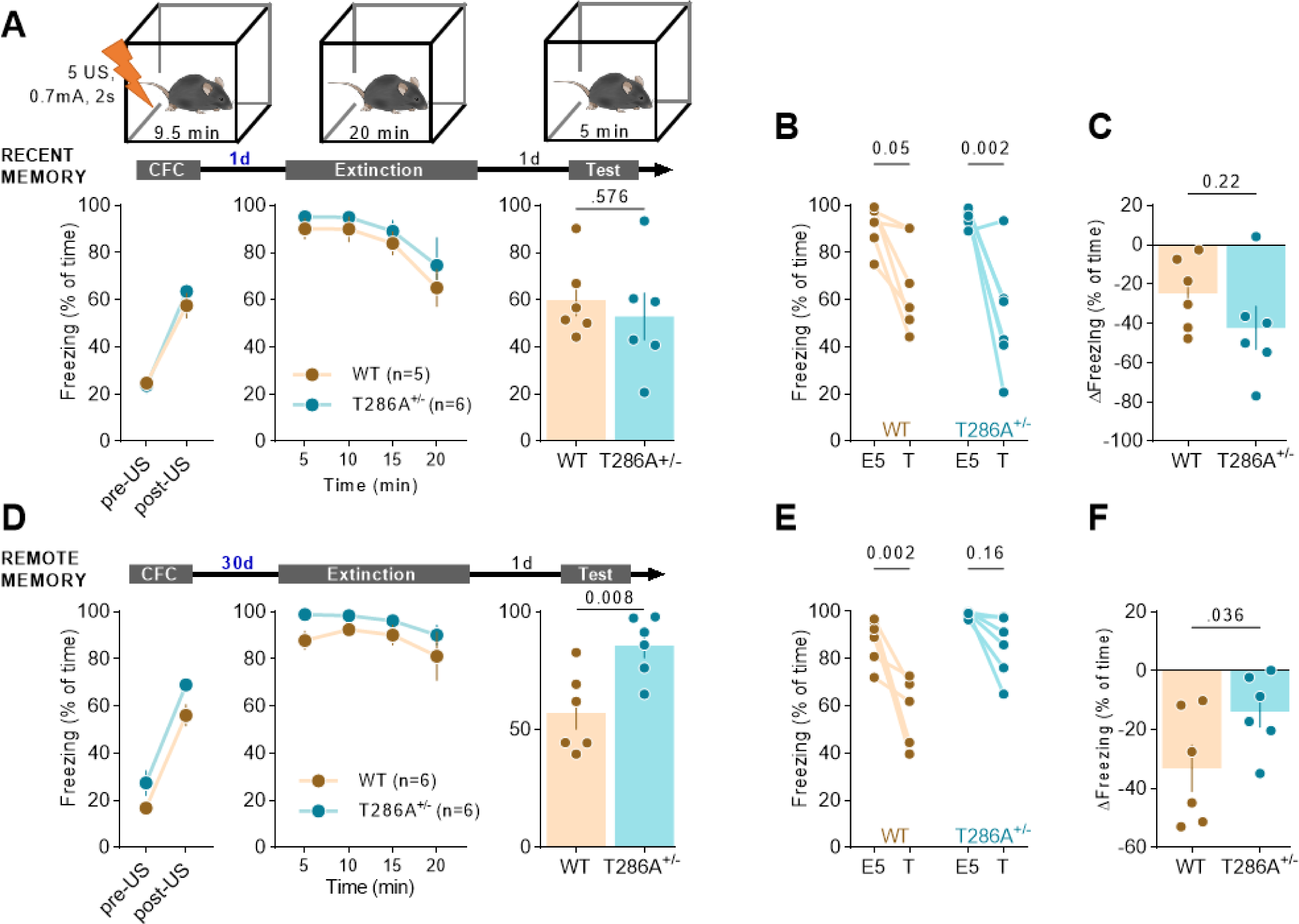
T286A^+/-^ mutation impairs extinction of remote, but not recent, contextual fear memory. **(A-C)** Experimental timeline and summary of data for freezing levels during the extinction of recent contextual fear memory. T286A^+/-^ and WT mice underwent CFC (CFC: RM ANOVA, effect of time: F(1, 10) = 236, P < 0.001; effect of genotype: F(1, 10) = 0.293, P = 0.600; time × genotype interaction: F(1, 10) = 2.20, P = 0.169), followed by recent fear memory extinction session 1 day later (Extinction: RM ANOVA, effect of time: F(1.24, 11.2) = 7.30, P = 0.016; effect of genotype: F(1, 9) = 1.25, P = 0.292; time × genotype interaction: F(3, 27) = 0.078, P = 0.971) and fear extinction test on the next day (Test: unpaired t-test: t(10) = 0.578, P = 0.576). **(B-C)** Summary of data showing freezing levels during the first 5 minutes of the extinction session (E5) and test (T) (RM ANOVA, effect of time: F(1, 10) = 25.33, P < 0.001; effect of genotype: F(1, 10) = 0.519, P = 0.49; time × genotype interaction: F(1, 10) = 1.728, P = 0.22), and the change of freezing frequency during T as compared to E5 (unpaired t-test: t(10) = 1.314, P = 0.22). **(D-F)** Experimental timeline and summary of data for freezing levels during the extinction of remote contextual fear memory. T286A^+/-^ and WT mice underwent CFC (CFC: RM ANOVA, effect of time: F(1, 10) = 186, P < 0.001; effect of genotype: F(1, 10) = 6.56, P = 0.028; time × genotype interaction: F(1, 10) = 0.143, P = 0.713), remote fear memory extinction session 30 days later (Extinction: RM ANOVA, effect of time: F(1.31, 13.1) = 2.06, P = 0.173; effect of genotype: F(1, 10) = 3.97, P = 0.074; time × genotype interaction: F(3, 30) = 0.170, P = 0.916) and fear extinction test on the next day (Test: unpaired t-test: t(10) = 3.271, P = 0.008). **(E-F)** Summary of data showing freezing levels during the first 5 minutes of the extinction session (E5) and test (T) (RM ANOVA, effect of time: F(1, 10) = 28.48. P < 0.001; effect of genotype: F(1, 10) = 21.81, P < 0.001; time × genotype interaction: F(1, 10) = 3.880, P = 0.08), and the change of freezing frequency during T as compared to E5 (unpaired t-test: t(10) = 2.01, P = 0.036).

Next, we tested the effect of T286A^+/-^ mutation on the extinction of remote contextual fear memory. This time the Extinction session was conducted 30 days after CFC. As previously, at the beginning of the Extinction session freezing levels were high in both experimental groups, indicating contextual fear memory recall, and slightly decreased during the session. Fear extinction memory was tested on the next day. While WT mice significantly decreased the frequency of freezing during the Test as compared to E5 indicating formation of fear extinction memory, such an effect was not observed for T286A^+/-^ mutants. The freezing levels of the heterozygotes during the Test did not significantly differ from the levels observed during E5, indicating no extinction of remote contextual fear memory (**Figure 1D-F**). Hence higher levels of autophosphorylated αCaMKII are required for updating of remote vs recent contextual fear memory.

### Neuronal correlates of remote contextual fear memory extinction

As T286A^+/-^ mutant mice showed impaired extinction of remote contextual fear memory we assumed that this mouse model is an excellent tool to study neuronal correlates of this phenomenon. To this end T286A^+/-^ mutant mice and their WT littermates underwent CFC and Extinction session 1 or 30 days after CFC (Recent and Remote groups, respectively). The animals were sacrificed 70 minutes after the Extinction sessions. In addition, we used 3 control groups: naive mice taken from home cages, 5US groups sacrificed 1 day after CFC without Extinction session, and context controls (Ctx) that were exposed to the experimental context twice but never received USs and were sacrificed 70 minutes after the last context exposure (**Supplementary Figure 1A**). Mice brains were sliced and used for immunofluorescent staining to detect c-Fos, as a marker of neuronal activity. c-Fos expression was analyzed in 23 brain regions of the dorsal hippocampus, amygdala, cortex, thalamus, and septum (**Supplementary Figure 1B**).

Overall, for both WT and T286A^+/-^ mutant mice, we observed a significant elevation of c-Fos levels in many brain regions following Recent and Remote extinction, as compared to 5US animals (**Figure 2**). The changes were, however, more pronounced after Remote as compared to Recent extinction. A significant increase in c-Fos levels was found in 12 regions after Recent, and 15 after Remote extinction in WT animals (**Figure 2A-B**); and in 12 regions after Recent, and 17 after Remote extinction in T286A^+/-^ mutants (**Figure 2C-D**). In five brain regions (RSG, MS, PV, BLA and LA) c-Fos levels were significantly higher in Remote vs Recent groups in WT animals, and in 10 regions (V1, ENT, LS, MS, AD, AV, CM, MD, PV, BLA) in heterozygotes. In addition, c-Fos levels were significantly lower in CA1 in the Remote vs Recent group in T286A^+/-^ mutant mice (**Figure 2C-D**). Furthermore, when we compared c-Fos levels between WT and T286A^+/-^ mutant mice the biggest differences were observed between the Remote extinction groups, as compared to all other experimental groups (**Figure 2E-F**). In the Remote groups, c-Fos levels were significantly higher in the heterozygotes in 9 regions (CG, V1, Ent, LS, MS, AD, RE, CM and BLA), and downregulated in DG. c-Fos levels were affected by the mutation in 5 regions in the naive groups, in 3 regions in Recent and Ctx groups and no significant differences were observed between the genotypes in 5US groups. Hence, elevated c-Fos levels in the cortex, septum, thalamus and amygdala, and reduced levels in the hippocampus are the neural correlates of impaired extinction of remote contextual fear memory in T286A^+/-^ mutant mice. This observation suggests that αCaMKII activity in these regions contributes to systems consolidation and updating of the remote memory engram.

**Figure 2.**
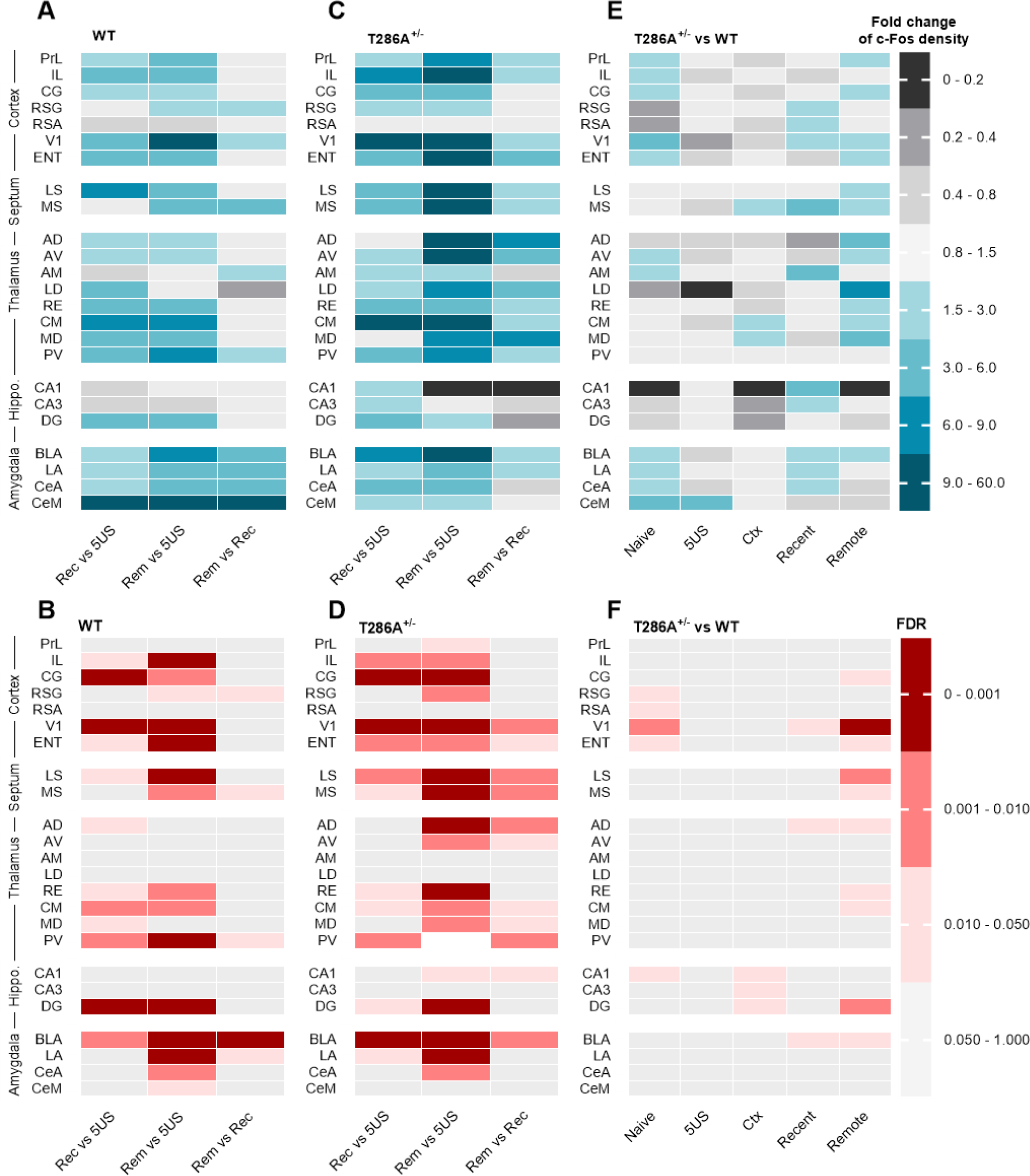
Summary of the c-Fos signal density profile comparison between groups for each structure within the brain region. **(A, C).** Fold change of the c-Fos signal density for WT and T286A+/- mice, where the recent (Rec) and remote (Rem) extinction groups are compared to mice sacrificed 24 hours after CFC (5US) or to each other. **(E)** Fold change of c-Fos signal density for T286A+/- mice as compared to WT animals. Naive, mice taken from home cages; Ctx, mice exposed twice to the novel context without US experience. See Supplementary Figure 1 for the experimental groups description. **(B, D, F)** Significant change of the c-Fos signal density for WT and T286A+/- mice (based on FDR results). PrL, prelimbic cortex; IL, infralimbic cortex; CG, cingulate cortex; RSG, retrosplenial cortex, granular part; RSA, retrosplenial cortex, agranular part; V1, primary visual cortex; ENT, entorhinal cortex; LS, lateral septum; MS, medial septum; AD, anterodorsal nucleus of thalamus; AV, anteroventral nucleus of thalamus; AM, anteromedial nucleus of thalamus; LD, laterodorsal nucleus of thalamus; RE, nucleus reuniens; CM, centromedial nucleus of thalamus; MD, mediodorsal nucleus of thalamus; PV, paraventricular nucleus of thalamus; CA1, dorsal hippocampus, CA1 area; CA3, dorsal hippocampus, CA3 area; DG, dentate gyrus; BLA, basolateral nucleus of the amygdala; LA, lateral nucleus of the amygdala; CeA, central nucleus of the amygdala, capsular and lateral divisions; CeM, central nucleus of the amygdala, centromedial division.

### Validation of the chemogenetic tools

As our c-Fos analysis revealed multiple brain regions that are likely regulated during extinction of remote contextual fear memory by αCaMKII, in the following experiment we validated their function in the process using a chemogenetic approach relying on the combination of Gi-protein-coupled inhibitory DREADDs (Designer Receptors Exclusively Activated by Designer Drugs) (hM4Di) and hM4Di agonist, clozapine N-oxide (CNO) ^30,31^. As a recent study showed that CNO is reverse-metabolized to clozapine and produces clozapine-like effects ^32^, we first tested the effect of CNO on remote fear memory extinction. C57BL/6J mice (without hM4Di expression) underwent CFC and 30 days later they were injected with CNO (0.5, 3 or 10 mg/kg) or saline. Thirty minutes later they were exposed to the training context for contextual fear memory extinction and fear extinction memory was tested in the same context one day later. We observed no effect of CNO on fear memory recall during the Extinction session, or fear extinction memory during the Test (**Supplementary Figure 2A-C**).

To test the efficiency of the chemogenetic manipulation to inhibit fear extinction-induced c-Fos expression, a new cohort of mice was injected into V1 with adeno-associated viral vectors (AAVs) encoding hM4Di and red fluorescent protein, mCherry, under human synapsin (*hSyn*) promoter allowing for neuron-specific transgene expression ^33^. As a control AAV encoding only mCherry was used. Three weeks post-surgery and viral transduction, mice underwent CFC, followed 30 days later by a contextual fear extinction session. Mice were injected with CNO (i.p., 3 mg/kg) 30 minutes prior to the Extinction and they were sacrificed 60 minutes after the Extinction session. Their brains were sliced and immunostained to detect c-Fos protein. The statistical analysis of the data revealed significantly lower frequency of c-Fos-positive nuclei among the cells transduced with hM4Di as compared to the mCherry controls indicating that chemogenetic manipulation effectively prevented c-Fos expression, hance blocked cell activity (**Supplementary Figure 2D-E**).

### Contribution of RE to extinction of contextual fear memory

Activity of RE is synchronized with freezing bouts [23,24], suggesting that activity patterns in the RE are related to suppression of fear responses. As c-Fos mapping identified that increased c-Fos expression in RE correlated with impaired extinction of remote contextual fear memory in the T286A^+/-^ mutant mice in the following experiment we tested the role of RE in extinction of recent and remote memory.

C57BL/6J mice were injected into RE with AAVs encoding hM4Di or mCherry (**Figure 3A-B**). Three (for recent groups) or one (for remote groups) week post-surgery and viral transduction mice underwent CFC, followed by a Recent or Remote contextual fear extinction session (1 or 30 days post CFC, respectively). Mice were injected with CNO (i.p., 3 mg/kg) prior to the Extinction. The freezing levels increased within the conditioning session (pre- vs post-US) and did not differ between the drug groups. During the Recent extinction session, the hM4Di mice had overall lower freezing levels as compared to the mCherry animals. Consolidation of recent fear extinction memory was tested 24 hours after the Extinction session in the same context (Test). Mice from the hM4Di group had lower levels of freezing as compared to the mCherry animals, suggesting facilitated extinction of recent fear memory (**Figure 3C**). However, further analysis of the behavioral data revealed that mice in both groups similarly decreased freezing levels in the Test, as compared to E5 (**Figure 3D-E**). Hence, lower freezing in hM4Di mice during the Test likely resulted from impaired recall of context fear memory during the Extinction session, rather than the effect on consolidation of fear extinction memory.

**Figure 3.**
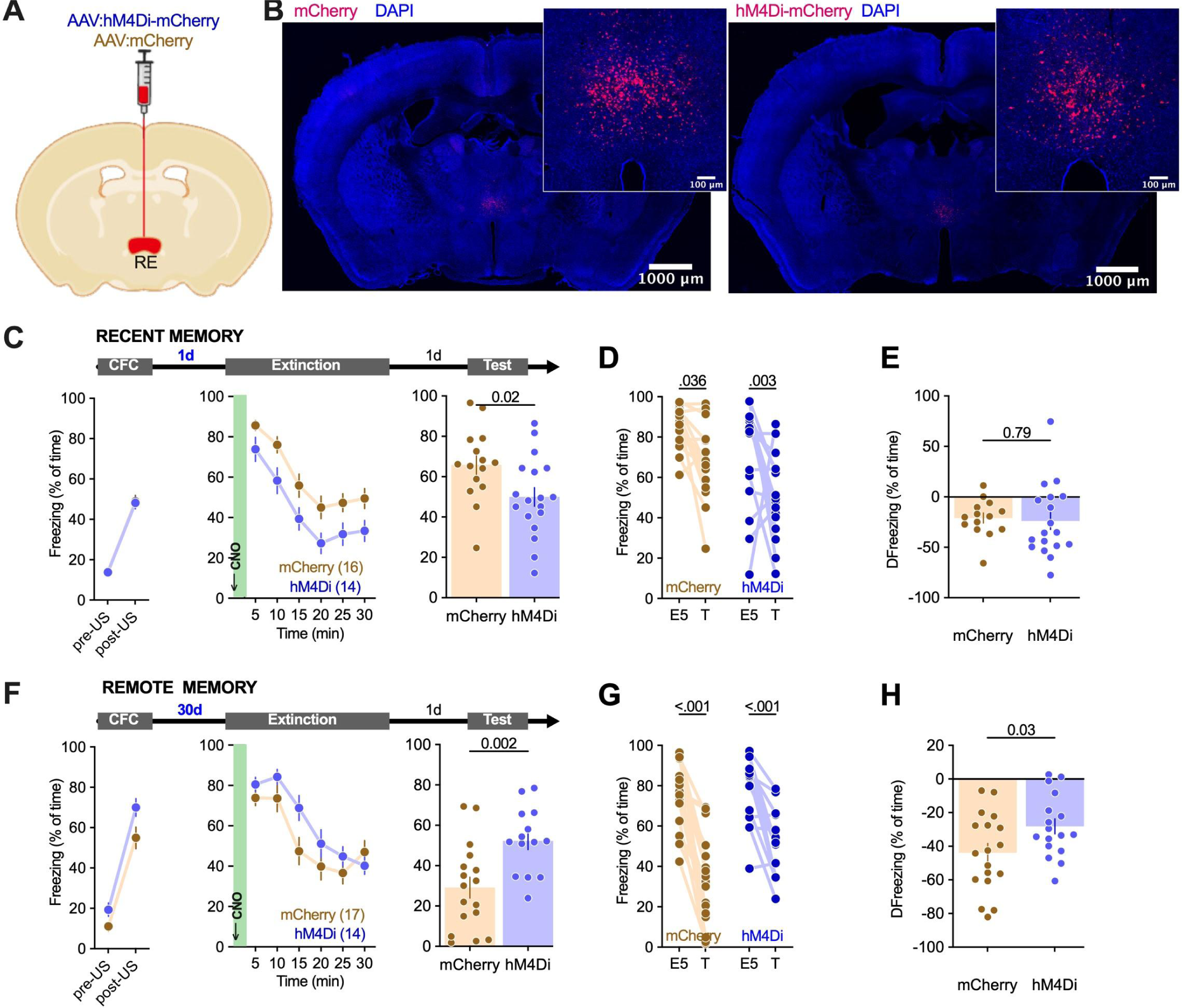
RE regulates extinction of remote contextual fear memory. **(A-B)** Surgery schema and representative microphotographs of AAV:hM4Di_mCherry (hM4Di) and AAV:mCherry (mCherry) expression in RE. **(C-E)** Experimental timeline and summary of data for freezing levels during the extinction of recent contextual fear memory. After stereotactic injection of hM4Di or mCherry into RE, mice underwent CFC (CFC: RM ANOVA, effect of time: F(1, 32) = 509, P < 0.001; effect of virus: F(1, 32) = 0.026, P = 0.871; time × virus interaction: F(1, 32) = 0.018, P = 0.892), followed by recent fear memory extinction session 1 day later (Extinction: RM ANOVA, effect of time: F(3.28, 78.8) = 39.2, P < 0.001; effect of virus: F(1, 24) = 2.05, P = 0.165; time × virus interaction: F(5, 120) = 2.16, P = 0.063) and fear extinction test on the next day (Test: unpaired t-test: t(25) = 1.63, P = 0.116). Mice were injected with CNO (i.p., 3 mg/kg) 30 minutes prior to the extinction session. **(D-E)** Summary of data showing freezing levels during the first 5 minutes of the extinction session (E5) and test (T) (RM ANOVA, effect of time: F(1, 26) = 43.9, P < 0.001; effect of virus: F(1, 26) = 2.69, P = 0.113; time × genotype interaction: F(1, 26) = 2.53, P = 0.124), and the change of freezing frequency during T as compared to E5 (unpaired t-test: t(26) = 1.59, P = 0.124). **(F-H)** Experimental timeline and summary of data for freezing levels during the extinction of remote contextual fear memory. After stereotactic injection of hM4Di or mCherry into RE, mice underwent CFC (CFC: RM ANOVA, effect of time: F(1, 31) = 443.8, P < 0.001; effect of virus: F(1, 31) = 3.680, P = 006; time × virus interaction: F(1, 31) = 1.226, P = 0.28), remote fear memory extinction session 30 days later (Extinction: RM ANOVA, effect of time: F(3.97, 115) = 27.8, P < 0.001; effect of virus: F(1, 29) = 2.64, P = 0.115; time × virus interaction: F(5, 145) = 1.98, P = 0.086) and fear extinction test on the next day (Test: unpaired t-test: t(29) = 3.337, P = 0.002). **(G-H)** Summary of data showing freezing levels during the first 5 minutes of the extinction session (E5) and test (T) (RM ANOVA, effect of time: F(1, 31) = 88.3, P < 0.001; effect of virus: F(1, 31) = 5.09, P = 0.031; time × virus interaction: F(1, 31) = 5.66, P = 0.024), and the change of freezing frequency during T as compared to E5 (unpaired t-test: t(33) = 2.201, P = 0.03).

During the Remote extinction session, mice in both experimental groups significantly decreased freezing levels indicating successful within session fear extinction, and no significant effect of the virus was observed. However, when the fear extinction memory was tested, the hM4Di animals demonstrated higher levels of freezing (**Figure 3F**). Further analysis of the behavioral data revealed that although both hM4Di and mCherry mice had lower levels of freezing during the Test as compared to E5, the change was larger in the control group (**Figure 3G-H**). Hence, chemogenetic inhibition of RE during the Remote, but not Recent, extinction session impaired consolidation of fear extinction memory.

### Contribution of MS to extinction of contextual fear memory

While, former studies showed that MS contributes to recent fear memory formation and extinction ^34–38^, our analysis indicates that MS activity during extinction increases as the memory age. To test the role of MS in extinction of remote fear memory C57BL/6J mice were injected into MS with AAVs encoding hM4Di or mCherry (**Figure 4A-B**). Three weeks post-surgery and viral transduction mice underwent CFC, followed by a Recent or Remote contextual fear extinction session (1 or 30 days post CFC, respectively). Mice were injected with CNO (i.p., 3 mg/kg) prior to the Extinction. The freezing levels increased during the CFC (pre- vs post-US) and did not differ between the drug groups. During the Recent extinction session, mice in both experimental groups decreased freezing levels and no significant effect of the virus was observed. Consolidation of recent fear extinction memory was tested 24 hours after the Extinction session in the same context (Test). Mice from the two experimental groups did not differ in freezing levels (**Figure 4C**). Both hM4Di and mCherry mice had lower levels of freezing during the Test as compared to E5, and the change was similar (**Figure 4D-E**), indicating that chemogenetic inhibition of MS during the Recent Extinction had no effect on the recall of contextual fear and consolidation of extinction memory.

**Figure 4.**
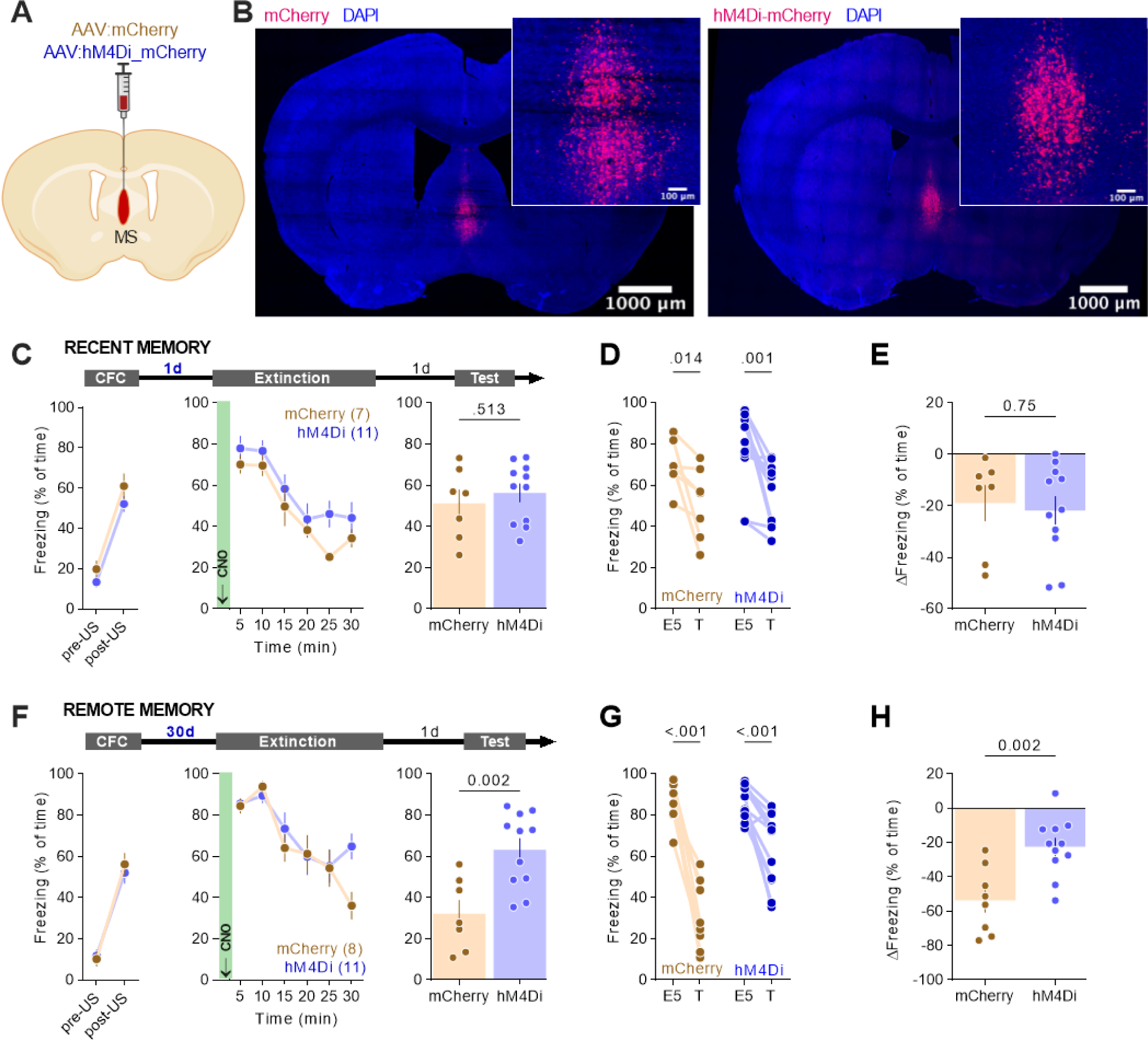
MS regulates extinction of remote contextual fear memory. **(A-B)** Surgery schema and representative microphotographs of AAV:mCherry (mCherry) and AAV:hM4Di_mCherry (hM4Di) expression in MS. **(C-E)** Experimental timeline and summary of data for freezing levels during the extinction of recent contextual fear memory. After stereotactic injection of hM4Di or mCherry into MS, mice underwent CFC (CFC: RM ANOVA, effect of time: F(1, 13) = 172, P < 0.001; effect of virus: F(1, 13) = 2.37, P = 0.148; time × virus interaction: F(1, 13) = 0.156, P = 0.699), followed by recent fear memory extinction session 1 day later (Extinction: RM ANOVA, effect of time: F(3.87, 62.0) = 26.3, P < 0.001; effect of virus: F(1, 16) = 1.93, P = 0.184; time × virus interaction: F(5, 80) = 0.689, P = 0.634) and fear extinction test on the next day (Test: unpaired t-test: t(16) = 0.669, P = 0.513). Mice were injected with CNO (i.p., 3 mg/kg) 30 minutes prior to the extinction session. **(D-E)** Summary of data showing freezing levels during the first 5 minutes of the extinction session (E5) and test (T) (RM ANOVA, effect of time: F(1, 16) = 21.5, P < 0.001; effect of virus: F(1, 16) = 0.968, P = 0.340; time × genotype interaction: F(1, 16) = 0.101, P = 0.755), and the change of freezing frequency during T as compared to E5 (unpaired t-test: t(16) = 0.669, P = 0.513). **(F-H)** Experimental timeline and summary of data for freezing levels during the extinction of remote contextual fear memory. After stereotactic injection of hM4Di or mCherry into MS, mice underwent CFC (CFC: RM ANOVA, effect of time: F(1, 17) = 193, P < 0.001; effect of virus: F(1, 17) = 0.061, P = 0.807; time × virus interaction: F(1, 17) = 0.864, P = 0.366), remote fear memory extinction session 30 days later (Extinction: RM ANOVA, effect of time: F(2.83, 48,0) = 18.3, P < 0.001; effect of virus: F(1, 17) = 0.819, P = 0.378; time × virus interaction: F(5, 85) = 2.42, P = 0.042) and fear extinction test on the next day (Test: unpaired t-test: t(16) = 3.585, P = 0.002). **(G-H)** Summary of data showing freezing levels during the first 5 minutes of the extinction session (E5) and test (T) (RM ANOVA, effect of time: F1, 17) = 82.6, P < 0.001; effect of virus: F(1, 17) = 11.7, P = 0.003; time × virus interaction: F(1, 17) = 13.9, P = 0.002), and the change of freezing frequency during T as compared to E5 (unpaired t-test: t(17) = 3.728, P = 0.002).

During the Remote extinction session, mice in both experimental groups significantly decreased freezing levels, indicating successful within session fear extinction, and no significant effect of the virus was observed. However, when the fear extinction memory was tested, the hM4Di animals demonstrated higher levels of freezing (**Figure 4F**). Further analysis of the behavioral data revealed that although both hM4Di and mCherry mice had lower levels of freezing during the Test as compared to E5, the change was significantly greater in the control group (**Figure 4G-H**). Hence, chemogenetic inhibition of MS during the Remote, but not Recent, extinction session impaired consolidation of contextual fear extinction memory.

### RE→MS projections

We hypothesized that projections between RE and MS contribute to neuronal networks that support extinction of remote contextual fear memory. As the data about the direct projections between RE and MS (RE→MS) are controversial ^39–43^, we first tested the presence of such projections using Ai14 mice that express Cre-dependent red fluorescent protein, tdTomato. Ai14 mice were stereotactically injected into MS with retro AAV (rAAV) encoding Cre recombinase allowing for transduction of axons terminating in MS and tdTomato expression in the cell bodies ^44,45^ (**Figure 5A-B**). tdTomato-positive cells were observed in the anterior (Bregma: 0.816 - 0.519 mm), but not the posterior part (Bregma: 0.473 to -0.586 mm) of RE (**Figure 5C**). We also observed tdTomato-positive cells in the brain regions well known to innervate MS, such as dCA1 and BLA ^46,47^ (**Figure 5C**). Similar results were observed when we injected rAAV encoding enhanced GFP (eGFP) to MS of C57BL/6J mice - eGFP-positive cells were found only in the anterior RE (data not shown). Hence, MS is innervated specifically by the anterior part of RE.

**Figure 5.**
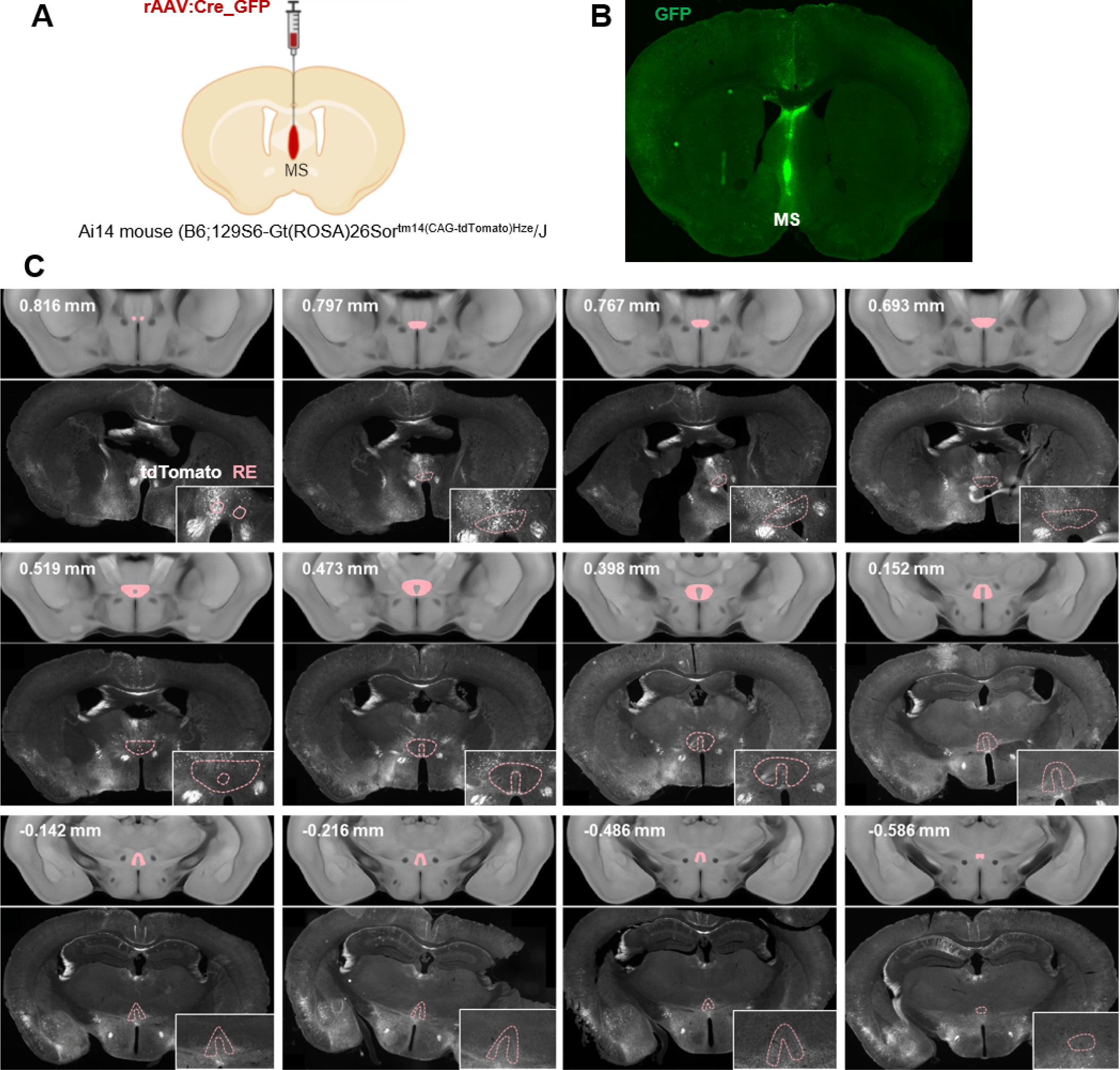
Identification of anatomical projections from anterior RE to MS. Ai14 mice (with Cre-dependent tdTomato) were injected into MS with rAAV:Cre_GFP resulting in tdTomato expression in neurons projecting to MS. **(A-B)** Surgery schema and representative microphotographs of rAAV:Cre_GFP expression in MS. **(C)** Microphotographs of tdTomato expression in Ai14 mouse (bottom) along with microphotographs of Allen mouse brain atlas (top). RE is marked in pink. Brain coordinates according to Bregma are indicated for each set of pictures.

### Contribution of RE→MS projections to extinction of contextual fear memory

To test whether RE→MS pathway affects extinction of contextual fear memory, C57BL/6J mice were stereotactically injected into RE with AAV encoding double-floxed inverted open reading frame (DIO) of hM4Di and mCherry, or just mCherry, under *camk2a* promoter (AAV:DIO_hM4Di_mCherry or AAV:DIO_mCherry), and into MS with rAAV encoding improved Cre recombinase (iCre) with eGFP under *camk2a* promoter (rAAV:iCre_eGFP). This combinatorial manipulation allowed for hM4Di expression specifically in RE neurons projecting to MS (RE→MS) (**Figure 6A-B**). Three weeks post-surgery and viral transduction, mice underwent CFC, followed by a Recent or Remote contextual fear extinction session. Mice were injected with CNO (i.p., 3 mg/kg) 30 minutes prior to the Extinction. The freezing levels increased within the CFC session (pre- vs post-US) in all experimental groups and did not differ between the virus groups. During the Recent Extinction session, the mice from two experimental groups had similar freezing levels at the beginning of the session and decreased freezing during the session, indicating no impairment of fear memory recall and successful within-session fear extinction. Consolidation of long-term fear extinction memory was tested 24 hours after the Extinction session in the same context (Test) (**Figure 6C**). Mice from two experimental groups had lower freezing levels during the Test as compared to E5, indicating successful consolidation of fear extinction memory (**Figure 6D**). No difference in freezing levels were observed between the experimental groups (**Figure 6E**).

**Figure 6.**
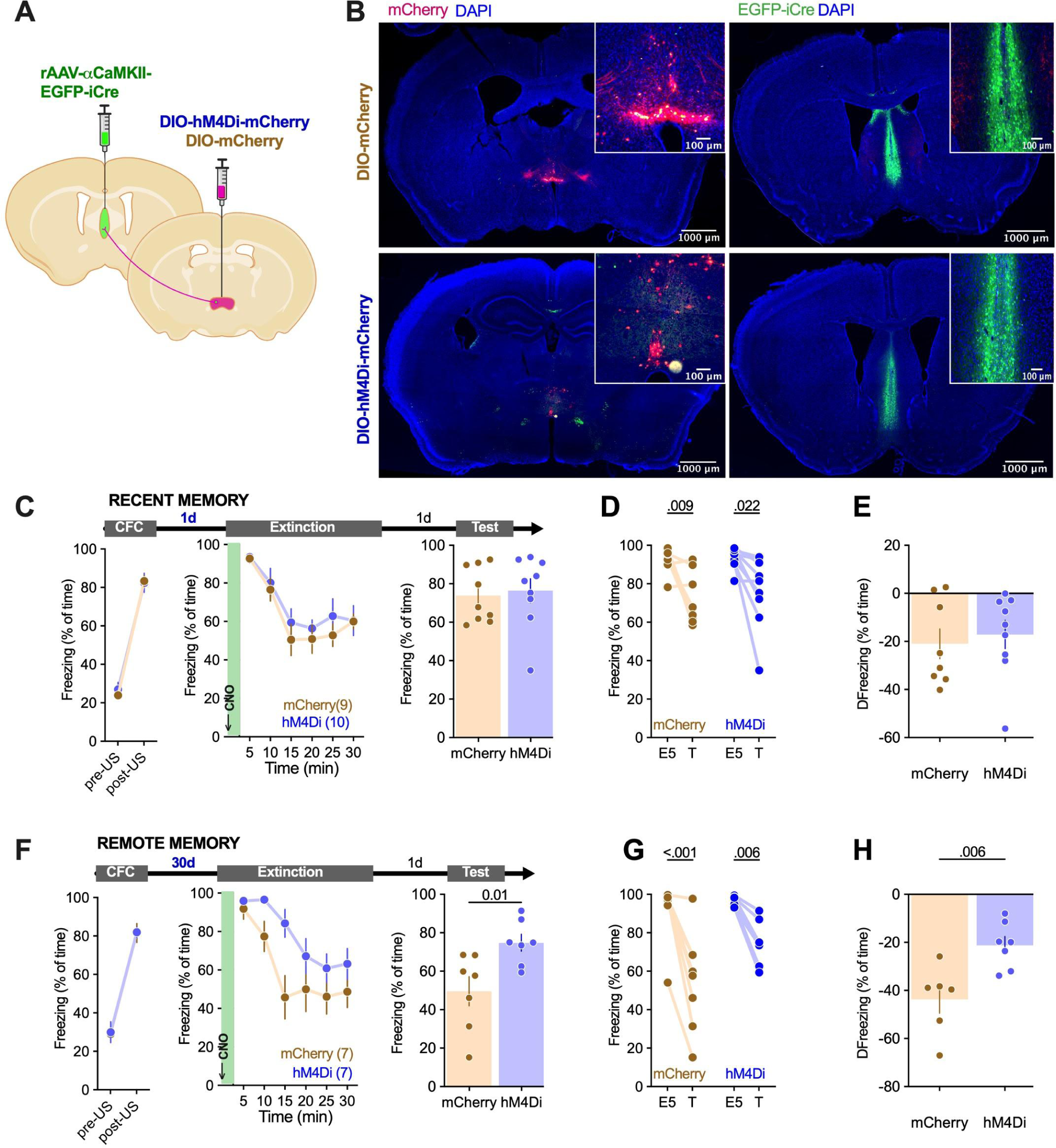
RE→ MS projections regulate consolidation of remote contextual fear extinction memory. AAV:DIO-hM4Di-mCherry (hM4Di) or control virus AAV:αCaMKII-DIO-mCherry (mCherry) were bilaterally injected to RE, while rAAV:iCre_eGFP-αCaMKII (eGFP) to MS of C57BL/6J male mice. **(A-B)** Surgery schema and representative microphotographs of hM4Di and mCherry in RE, and eGFP expression in MS. **(C-E)** Experimental timeline and summary of data for freezing levels during the extinction of recent contextual fear memory. After stereotactic injection of hM4Di or mCherry into RE and eGFP into MS, mice underwent CFC (CFC: RM ANOVA, effect of time: F(1, 15) = 414, P < 0.001; effect of virus: F(1, 15) = 0.056, P = 0.82), followed by recent fear memory extinction session 1 day later (Extinction: RM ANOVA, effect of time: F(1, 15) = 19.68, P < 0.001; effect of virus: F(1, 15) = 0.648, P = 0.43) and fear extinction test on the next day (Test: unpaired t-test: t(16) = 0.318, P = 0.75). Mice were injected with CNO (i.p., 3 mg/kg) 30 minutes prior to the extinction session. **(D-E)** Summary of data showing freezing levels during the first 5 minutes of the extinction session (E5) and test (T) (RM ANOVA, effect of time: F(1, 15) = 19.7, P < 0.001; effect of virus: F(1, 15) = 0.405, P = 0.534; time × virus interaction: F(1, 15) = 0.201, P = 0.660), and the change of freezing frequency during T as compared to E5 (unpaired t-test: t(15) = 0.448, P = 0.66). **(F-H)** Experimental timeline and summary of data for freezing levels during the extinction of remote contextual fear memory. After stereotactic injection of hM4Di or mCherry into MS, mice underwent CFC (CFC: RM ANOVA, effect of time: F(1, 11) = 201.3, P < 0.001; effect of virus: F(1, 11) = 0.0190, P = 0.89; time × virus interaction: F(1, 11) = 0.0057, P = 0.94), remote fear memory extinction session 30 days later (Extinction: RM ANOVA, effect of time: F(5, 65) = 13.60, P < 0.001; effect of virus: F(1, 13) = 5.402, P = 0.04; time × virus interaction: F(5, 65) = 1.465, P = 0.21) and fear extinction test on the next day (Test: unpaired t-test: t(12) = 2.88 P = 0.01). **(G-H)** Summary of data showing freezing levels during the first 5 minutes of the extinction session (E5) and test (T) (RM ANOVA, effect of time: F(1, 13) = 52.1, P < 0.001; effect of virus: F(1, 13) = 2.33, P = 0.151; time × virus interaction: F(1, 13) = 3.77, P = 0.074), and the change of freezing frequency during T as compared to E5 (unpaired t-test: t(11) = 3.39, P = 0.006).

During the Remote extinction session, mice in both experimental groups significantly decreased freezing levels indicating successful within session fear extinction. The hM4Di groups showed, however, overall higher freezing levels. Moreover, the hM4Di animals also demonstrated higher levels of freezing when the fear extinction memory was tested (**Figure 6F**). Further analysis of the behavioral data revealed that although both hM4Di and mCherry mice had lower levels of freezing during the Test as compared to E5, the change was significantly greater in the control group (**Figure 6G-H**). Hence, chemogenetic inhibition of RE→MS projection during the Remote, but not Recent, extinction session impaired consolidation of fear extinction memory.

## Discussion

The current study identifies neuronal networks involved in the extinction of remote contextual fear memory. In particular, we characterize the role of RE, MS, and RE→MS projections in the extinction of contextual fear memory as the memories age.

In our experiments, we evaluated extinction of contextual fear memory at both short and long delays after training. We found that in the wild-type (WT) animals functional networks differed at these two time-points, consistent with the idea that memory organization changes as a function of memory age, a process known as systems consolidation ^12^. One prominent model of systems consolidation ^48^ proposes that memories are initially encoded in hippocampal-cortical networks, and that reactivation of these networks then leads to the incremental strengthening of inter-cortical connectivity, and an emergent role for cortical regions in memory expression coupled with disengagement of the hippocampus. In agreement with this model we see increased activity of the cortex (RSC and V1) in the remote time point. Furthermore, we see increased engagement of the septum (MS), thalamus (PV) and amygdala (BLA and LA) supporting the contribution of these regions in systems consolidation ^15,29^. However, we do not see significant change in the hippocampus engagement with time, possibly indicating differences between the contribution of this region in contextual fear memory formation, recall and extinction as the memory age. Interestingly, the role of some hotspots, including BLA, RSC, PV and PV→CeA projections, in remote memory processing has been already validated ^29,49,50^.

Overall, our data indicates that extinction of remote memory engages more brain regions than recent memory. Interestingly, although previous studies linked broader memory networks with higher stabilization of remote memory ^15,51,52^, in our experiments extinction of remote memory was more efficient. A remote extinction session resulted in WT mice in a reduction of freezing by 40-50% during the extinction test as compared to the beginning of the extinction session, while the recent extinction session reduced freezing by 20% measured during the test. Hence, increased stabilization of memories with time may be specific for traumatic and pathologic memories ^53,54^, while in case of physiologic process extinction of remote fear memory may be enhanced by natural forgetting of aversive experience. Alternatively, enhanced extinction observed in our model in the remote time point may reflect the fact that such a session enggages more brain regions, hence, more efficiently remodels memory networks.

We show that αCaMKII autophosphorylation deficient heterozygous mice (T286A^+/-^) have impaired extinction of remote memory while the extinction of recent contextual fear memory is spared. Accordingly, we use this model to look for neuronal correlates of remote memory extinction. Although previous studies showed that T286A mutation affects extinction of recent contextual fear memory (more training trials are required for efficient extinction) ^20–22^, this is the first demonstration showing that αCaMKII autophosphorylation affects remote memory updating. This finding is reminiscent of former studies showing that αCaMKII^+/-^ mice show impairment of remote memory storage ^15,19^. Impaired remote memory extinction in T286A^+/-^ mice is accompanied with globally increased activity of the thalamus (RE, CM, AD), septum (LS and MS), cortex (CG, V1, ENT) and amygdala (BLA), and decreased activity of the hippocampus (DG). Hence, impaired fear memory extinction in heterozygous may result from aberrant connectivity of fear extinction network and impaired systems consolidation. It is also possible that the decreased function of the hippocampus in the mutants during the Remote extinction session reflects impaired recall and/or updating of contextual fear memory.

While correspondence between the predictions derived from the c-Fos analysis and these published findings provides some validation of our approach, perhaps greater value lies in using this approach for the identification of novel, candidate brain regions. In this regard, it is interesting that both septal (MS and LS) and thalamic (RE, CM, AD) regions were identified. These regions, by virtue of their widespread connectivity with both the hippocampus and neocortex, are ideally positioned to influence network function. In particular, thalamic regions strongly influence cortical activity and therefore may play central roles in coordinating memory retrieval and extinction.

Indeed, former studies demonstrated the role of the RE in fear memory encoding, retrieval, extinction and fear generalization ^55–60^. Specifically, it has been demonstrated that RE is involved in the extinction of recent contextual and auditory fear memory ^61,62^. Administration of muscimol into the RE before the recent extinction session impairs whitin-session fear extinction as well as fear extinction memory tested on the following day ^61,62^. Administration of muscimol into the RE shortly after extinction training or extinction reconsolidation does not result in differences in freezing levels during the extinction memory test ^63^. In opposition to these findings our experiments indicate that the involvement of the RE in fear memory extinction depends on the length of the interval between training and extinction sessions. Chemogenetic inhibition of RE neurons during the extinction of recent contextual fear memory slightly increased freezing levels during the extinction session but did not affect the consolidation of extinction memory. Chemogenetic inhibition of RE neurons during the extinction of remote contextual fear memory impaired both within-session extinction and consolidation of extinction memory. Reminiscent of these observations, permanent lesion of the RE impaired remote, but not recent, contextual fear memory ^64^ and chemogenetic manipulation of RE bidirectionally affects extinction of the remote contextual fear memory ^29^. The discrepancy between our results and the work of Marren’s group ^63^ may stem from differences in the location of RE inhibition along the anteroposterior axis and the methods used to inhibit RE activity as well as training protocol. From the above work, it can be inferred that the researchers primarily inhibited the posterior part of RE, whereas in our experiments, mainly the anterior part of RE was manipulated. Anatomical studies of RE indicate that its posterior part innervates mPFC, while the anterior RE innervates CA1 and ENT ^65–67^. In addition, our work demonstrates projections of the anterior RE to MS that are specifically involved in the remote extinction. Furthermore, muscimol used by Ramanathan et al. ^58^ stimulates inhibitory GABAA receptors and strongly inhibits neuronal activity. Whereas, stimulation of the inhibitory hM4Di receptor results in a decrease rather than complete inhibition of neuronal activity ^68^. Finally, in the former studies, as compared to our experiments, the authors used a weaker training protocol (3US vs 5US) and extended extinction session (35 min vs 30 min) that resulted in more efficient contextual fear extinction (to 20% vs 50-60% freezing). Hence, high levels of freezing observed in our experiments in the control groups following a recent extinction session could mask the effect of RE inhibition on contextual fear extinction. In summary, our experiments suggest that the role of RE in extinction of contextual fear memory increases with time. Moreover, the extinction of remote and recent fear memory possibly involves different populations of RE neurons. The later hypothesis should be, however, validated in the future experiments.

Several reports demonstrate the involvement of MS in extinction of fear memory ^38,69,70^. In conjunction with anatomical data indicating dense connections of MS with the hippocampus ^71–74^ and the medial prefrontal cortex ^75^, the above observations suggest the involvement of MS in processing spatial information related to environmental threats. In particular, cholinergic neurons of MS regulate the extinction of recent contextual fear memory when extinction training consists of seven sessions conducted over seven days ^76^. Lesions of MS and the adjacent nucleus of the diagonal band of Broca also led to impaired extinction of early fear memory dependent on auditory stimuli in rats ^77^. Here, we observed that c-Fos protein levels in MS are higher after the remote extinction session than after the recent extinction. Moreover, we did not observe a disruption of the extinction with chemogenetic inhibition of MS during the recent extinction session. However, we noticed that chemogenetic inactivation of MS during the remote extinction session impaired formation of the fear extinction memory. Thus, the changes in c-Fos expression and the results of chemogenetic manipulations are consistent and suggest that the involvement of MS in fear memory extinction changes as the memory age. These observations, however, seem to contradict the literature data cited above ^76^. Possible reasons for this discrepancy may include differences in the conditioning and fear memory extinction protocols, as well as in the method and extent of MS neuron inactivation. Although Tronson et al. ^76^ observed impaired recent fear memory extinction in mice with MS lesions compared to the control animals, these differences were only evident on the fourth day of extinction training. This observation is similar to the results of our experiment. Interestingly, this difference probably does not depend on the length of the extinction session (3 min in Tronson et al. ^76^ vs. 30 min here), as in the study by Knox and Keller ^36^, which used long extinction training, a similar pattern of changes in behavior was observed as after 3-minute extinctions. Therefore, the time from the conditioning to the extinction session seems to be crucial for the involvement of MS in fear memory extinction.

RE innervates many brain structures and is innervated by over 30 areas ^39^. It also has about 11% bifurcating neurons, which innervate both the cingulate cortex and the medial part of the prefrontal cortex, as well as the insular cortex ^78^. So far, the functionally best-understood neuronal circuit involving RE is the mPFC→RE→hippocampal axis. Both mPFC→RE and RE→CA1 projections mediate the fear extinction process ^26,27,61^. It has been shown that inhibiting RE→CA1 projections during the recall of recent fear memory leads to impaired fear memory extinction in subsequent days ^79^. Furthermore, RE→BLA projection participates in the extinction of remote contextual fear memory [16]. It has been shown that both the activity of excitatory RE→BLA projections and the activity of RE cells increase shortly before the end of the extinction episode. Moreover, chemogenetic inhibition of these projections during late fear extinction impairs the extinction of late fear memory. On the other hand, optogenetic stimulation of GABAergic or dopaminergic cells in the zone incerta (ZI)→RE neuronal pathway leads to enhanced extinction of early fear memory dependent on auditory stimuli ^80^, revealing a complex role of RE in the neuronal circuits of fear extinction. Here, we demonstrated that chemogenetic inhibition of the excitatory RE→MS projections during fear memory extinction impairs extinction of remote, but not recent, contextual fear memory. Considering the consequences of inhibiting RE→MS projections in the context of anatomical connections, it is worth mentioning that both these regions innervate the CA1 area of the hippocampus, which is involved in extinction of contextual fear memory ^28,81–84^. MS and RE synchronize theta rhythms in the hippocampus and influence local field potentials in the hippocampus→RE→mPFC circuit ^85,86^.

In summary, the results presented here reveal a new element of neuronal circuits responsible for the extinction of remote contextual fear memory and highlight the importance of thalamo-septal circuits in the systems consolidation of remote memories.

## Materials and Methods

### Animals

C57BL/6J male mice were purchased from the Medical University of Białystok, Poland. T286A^+/-^ and WT mice were obtained from crossing heterozygous αCaMKII autophosphorylation-deficient mutant mice (αCaMKII-T286A^+/-^) ^23^ and heterozygotes of Thy1-GFP(M) line (Thy1-GFP^+/-^) ^87^ at the Nencki Institute animal house, and genotyped as previously described ^23,87^. Homozygous Ai14 mice were genotyped as previously described ^45^ and crossed with C57BL/6J animals. Heterozygotes were used in the experiment. Mice were 10 weeks-old at the beginning of the training. The animals were maintained on a 12 h light/dark cycle with food and water *ad libitum*. Behavioral experiments were conducted in the light phase of the cycle. All procedures conducted in the study were approved by the 1st Local Ethical Committee, Warsaw, Poland (permission number: 261/2012 and 529/2018).

### Contextual fear conditioning (CFC)

Mice were trained in a conditioning chamber located in a soundproof box (Med Associates Inc, St Albans, VT, USA). The chamber floor had a stainless steel grid for shock delivery. Before and after each training session, chambers were cleaned with 70% ethanol, and paper towels soaked in ethanol were placed under the grid floor. A video camera was fixed inside the door of the sound attenuating box, for the behavior to be recorded. Freezing behavior (defined as complete lack of movement, except respiration) and locomotor activity of mice were automatically scored. The experimenters were blind to the experimental groups.

On the day of CFC, mice were brought to the conditioning room 30 minutes before the training. Animals were placed in the chamber, and after a 148-second introductory period, a foot shock (2 seconds, 0.70 mA) (US) was presented. The shock was repeated five times, with intertrial interval of 90 seconds. Thirty seconds after the last shock, mice were returned to the home cage. Mice of the 5US groups were sacrificed 24 hours after the initial training. For the extinction of contextual fear, mice were re-exposing to the conditioning chamber without US presentation for 20 (WT and T286A^+/-^ animals) or 30 minutes (C57BL/6J mice) either 1 or 30 days after the training (extinction of recent and remote memory, respectively). WT and T286A^+/-^ mice from the Extinction groups were sacrificed 70 minutes after the extinction session. To test the efficiency of extinction, mice were re-exposed to the training context 24 hours after the extinction session for an additional 5 minutes. The differences in the length of the extinction session used for T286A^+/-^ animals and C57BL/6J mice resulted from the fact that while for the WT/T286A^+/-^ mice a 20 minute-long extinction session efficiently decreased freezing during the test, we did not see such effect for the C57BL/6J mice. Accordingly, we had to adjust the extinction protocol for these mice.

#### CNO administration

For intraperitoneal (i.p.) injections, Clozapine N-Oxide (CNO) was dissolved in 0.9% saline. If not stated otherwise, CNO (3 mg/kg) was injected 30 minutes prior to the contextual fear memory extinction session.

### c-Fos immunoreactivity

The mice were anesthetized with a mixture of Ketamine/Xylazine (90 mg/kg and 7,5 mg/kg respectively, (i.p.) Biowet Puławy) and next sedated with sodium pentobarbital (50 mg/kg intraperitoneal (i.p.) and perfused transcardially with 20 ml of PBS (Phosphate-buffered saline; POCH, Poland), followed by 50 ml of 4% paraformaldehyde (PFA; Sigma-Aldrich, Poland) in 0.1 M PBS, pH 7.5. Brains were stored in the same solution for 24 h at 4℃. Next, the brains were transferred to the solution of 30% sucrose in 0.1 M PBS (pH 7.5) and stored at 4℃ for 72 h. Brains were frozen at -20℃ and coronal sections (40 μm) cut in the frontal plane on a cryostat microtome (YD-1900, Leica). Coronal slices were stored in -20℃ in PBSAF (PBS, 15% sucrose, 30% ethylene glycol, 0.05% NaN_3_). Every sixth section through the whole brain was used for immunostaining. Sections were washed three times for 6 min in PBS, followed by incubation in blocking solution (5% NDS; Jackson Immuno Research/0,3% Triton X-100; Sigma Aldrich) in PBS for 2 h at room temperature (RT). Next, sections were incubated with c-Fos polyclonal antibody (sc-52; Santa Cruz Biotechnology, 1:1000) for 12 h in 37℃, washed three times in TBS (0,3% Triton X-100 in PBS) and incubated with secondary antibody (1:500) conjugated with Alexa Fluor-594 (Invitrogen A-11062) for 2 h at room temperature. After incubation with the secondary antibody sections were washed three times for 6 min in PBS and mounted on microscopic slides covered with mounting dye with DAPI (Invitrogen, 00-4959-52).

#### c-Fos quantification

c-Fos immunostaining was imaged under a confocal microscope (Leica TCS SP5; objective 10x). Cells were considered positive for c-Fos immunoreactivity if the nucleus had an area ranging from 5 to 160 μm^2^, and was significantly distinct from the background. c-Fos-positive cells were counted in 23 brain regions. We took 2 to 8 microphotographs for each animal per each brain region, resulting in 9 to 48 pictures per experimental group per brain region. Microphotographs were converted into an 8-bit grayscale and the same threshold used for a selected area of interest. The densities of the nuclei were counted with the ImageJ software (National Institute of Health, Bethesda, MD) by an experimenter blind to the experimental condition. The position of the analyzed brain regions was determined according to the mouse brain atlas ^88^ (**Supplementary Figure 1B-C**).

### Stereotactic surgery

Mice were anesthetized with isoflurane (5% for induction, 1.5-2.0% after) and fixed in the stereotactic frame (51503, Stoelting, Wood Dale, IL, USA). AAV_2.1_ viruses [camk2a_mCherry (titer 7.5 × 10^7^ pc/μl, from Karl Deisseroth’s Lab, Addgene plasmid #114469); hSyn_hM4D(Gi)_mCherry (4.59 × 10^9^ pc/ul; Addgene plasmid #50475); were injected into RE (250 nl, coordinates from the Bregma: AP, -0.58 mm; ML, ±0,1 mm; DV, -4.3 mm), MS (150 nl, coordinates from the Bregma: AP, -0,75 mm; ML, ± 0.0 mm; DV, -4.1 mm), hSyn_DiO_hM4Di-mCherry (3.3 × 10^9^ pc/µl; constructed by the VVF [dlox-hM4D(Gi)_mCherry(rev)-dlox: Addgene #50461]) and hSyn_DIO_mCherry (3.4 × 10^9^ pc/µl; constructed by the VVF) and camk2a_hM4D(Gi)_mCherry (4.4*10^9^ pc/ul; Addgene plasmid #50477) were injected into RE; rAAV:iCre_eGFP (3.6 × 10^9^ pc/µl) constructed by the VVF [iCre: Addgene #24593] was injected into the MS ^88^. We used beveled 26 gauge metal needle and 10 µl microsyringe (SGE010RNS, WPI, USA) connected to a microsyringe pump (UMP3, WPI, Sarasota, USA), and its controller (Micro4, WPI, Sarasota, USA) with an injection rate 0.1 µl/min. After injection, the needle was left in place for an additional 10 min to prevent unwanted spread of the vector. The AAVs were prepared by the Laboratory of Animal Models at Nencki Institute of Experimental Biology, Polish Academy of Sciences and Viral Vector Facility of ETH Zurich. Efficient and specific viral expression was assessed on the brain sections after all experiments. Only mice with AAVs in the regions of interest were used in the statistical analyses.

### Statistics

Analysis was performed using Graphpad Prism 10. All the statistical details of experiments can be found in the legends of the figures. Data with normal distribution are presented as mean ± standard error of the mean (SEM) or as median ± interquartile range (IQR) for the population with non-normal distribution. When the data met the assumptions of parametric statistical tests, results were analyzed by one- or repeated measures two-way ANOVA (RM ANOVA), followed by Tukey’s or Fisher’s *post hoc* tests, where applicable. Differences in means were considered statistically significant at p < 0.05.

## Acknowledgments, Funding and Disclosure

This work was supported by a National Science Center (Poland) [PRELUDIUM Grant No. 2019/35/N/NZ4/01910 to KFT; and MAESTRO grant No. 2020/38/A/NZ4/00483 to KR]. The project was carried out with the use of CePT infrastructure financed by the European Union - The European Regional Development Fund within the Operational Program “Innovative economy” for 2007-2013.

The funders had no role in study design, data collection and analysis, decision to publish, or preparation of the manuscript.

## DECLARATION OF INTEREST

Authors report no financial interests or conflicts of interest.

## INCLUSION AND DIVERSITY

While citing references scientifically relevant for this work, we also actively worked to promote gender balance in our reference list. We support inclusive, diverse, and equitable conduct of research.

## Supplementary materials

**Supplementary Figure 1.**
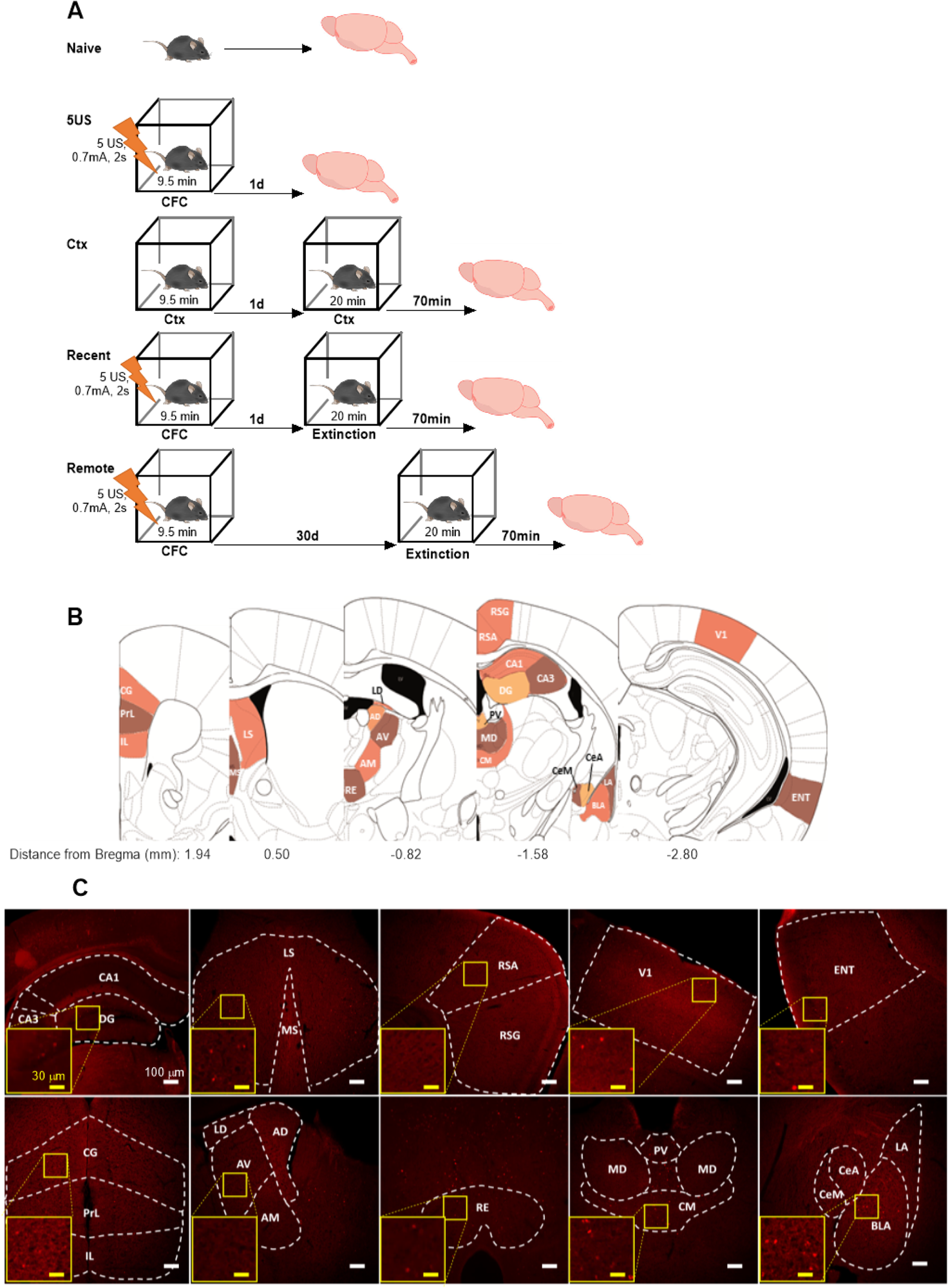
Experimental design for c-Fos analysis. **(A)** Experimental groups. Naive mice were taken from their home cage. 5US group underwent CFC with 5 electric shocks in novel context and were sacrificed 1 day later. Ctx mice were exposed twice to the novel context without US experience and were sacrificed 70 minutes after the last exposure. Recent groups underwent CFC and a contextual fear extinction session 1 day after CFC, and were sacrificed 70 minutes after the extinction. Remote groups underwent CFC and a contextual fear extinction session 30 days after CFC, and were sacrificed 70 minutes after the extinction. Mice brains were sliced and used for c-Fos immunostaining. **(B)** Analised brain regions. PrL, prelimbic cortex; IL, infralimbic cortex; CG, cingulate cortex; RSG, retrosplenial cortex, granular part; RSA, retrosplenial cortex, agranular part; V1, primary visual cortex; ENT, entorhinal cortex; LS, lateral septum; MS, medial septum; AD, anterodorsal nucleus of thalamus; AV, anteroventral nucleus of thalamus; AM, anteromedial nucleus of thalamus; LD, laterodorsal nucleus of thalamus; RE, nucleus reuniens; CM, centromedial nucleus of thalamus; MD, mediodorsal nucleus of thalamus; PV, paraventricular nucleus of thalamus; CA1, dorsal hippocampus, CA1 area; CA3, dorsal hippocampus, CA3 area; DG, dentate gyrus; BLA, basolateral nucleus of the amygdala; LA, lateral nucleus of the amygdala; CeA, central nucleus of the amygdala, capsular and lateral divisions; CeM, central nucleus of the amygdala, centromedial division. **(C)** Representative microphotographs of c-Fos immunostaining in the analyzed brain regions.

**Supplementary Figure 2.**
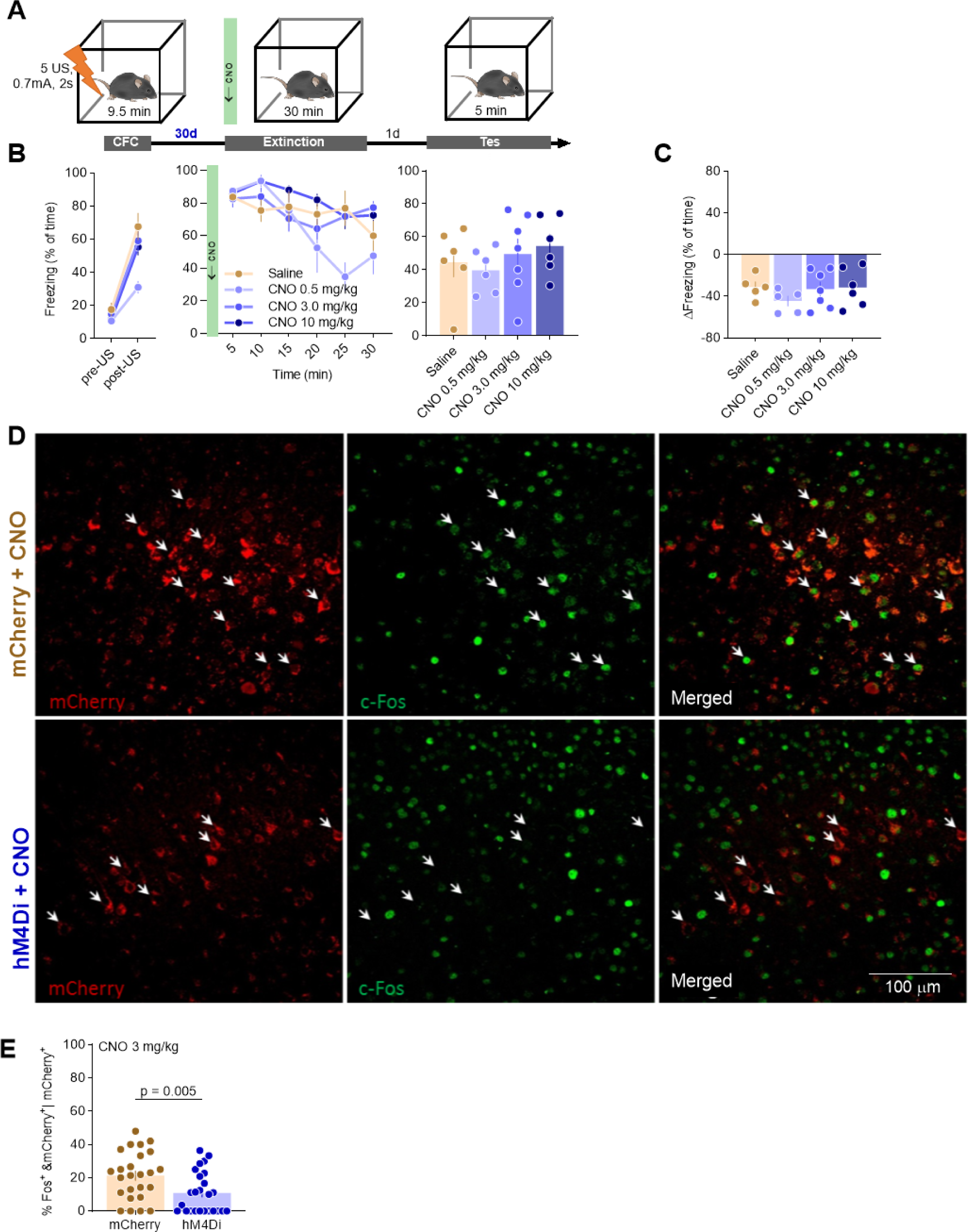
Validation of chemogenetic tools. **(A)** Experimental timeline. C57BL/6J mice (without hM4Di expression) underwent CFC and 30 days later they were injected with CNO (0.5, 3 or 10 mg/kg) or saline. Thirty minutes later they were exposed to the training context for contextual fear memory extinction and fear extinction memory was tested in the same context one day later. **(B)** Summary of data for freezing levels during CFC (CFC: RM ANOVA, effect of time: F(1, 21) = 270, P < 0.001), remote fear memory extinction session (Extinction: RM ANOVA, effect of time: F(3.66, 69.5) = 11.2, P < 0.001; effect of treatment: F(3, 19) = 2.59, P = 0.083) and fear extinction test (Test: one-way ANOVA, F(3, 21) = 0.613, P = 0.614). **(C)** Summary of data showing the change of freezing frequency during Test (T) as compared to the first 5 minutes of the Extinction (E5) (unpaired t-test: one-way ANOVA, F(3, 19) = 0.846, P = 0.486). **(D-E)** Analysis of the efficiency of chemogenetic inhibition. **(D)** Representative microphotographs of c-Fos immunostaining and hM4Di and mCherry expression in V1. **(E)** Summary of data analysis for the frequency of c-Fos-positive cells among the cells transduced with hM4Di or mCherry (Mann-Whitney U = 183, P = 0.005).

## Notes

### Competing Interest Statement

The authors have declared no competing interest.

